# Microbiome-derived metabolites reproduce the mitochondrial dysfunction and decreased insulin sensitivity observed in type 2 diabetes

**DOI:** 10.1101/2020.08.02.232447

**Authors:** Michael J. Ormsby, Heather Hulme, Victor H. Villar, Gregory Hamm, Giovanny Rodriguez-Blanco, Ryan A. Bragg, Nicole Strittmatter, Christopher J. Schofield, Christian Delles, Ian P. Salt, Saverio Tardito, Richard Burchmore, Richard J. A. Goodwin, Daniel M. Wall

**Affiliations:** Institute of Infection, Immunity and Inflammation, College of Medical, Veterinary and Life Sciences, Sir Graeme Davies Building, University of Glasgow, Glasgow G12 8TA, United Kingdom; Imaging and data Analytics, Clinical Pharmacology and Safety Sciences, R&D, AstraZeneca, Cambridge CB4 0WG, United Kingdom; Cancer Research UK Beatson Institute, Garscube Estate, Switchback Road, Glasgow G61 1BD, United Kingdom; Pharmaceutical Sciences, BioPharmaceuticals R&D, AstraZeneca, Cambridge CB4 0WG, United Kingdom; Chemistry Research Laboratory, University of Oxford, Mansfield Road, Oxford OX1 3TA, United Kingdom; Institute of Cardiovascular and Medical Sciences, College of Medical, Veterinary and Life Sciences, University of Glasgow, Glasgow G12 8TA, United Kingdom; Institute of Cancer Sciences, University of Glasgow, Glasgow G61 1QH, United Kingdom

## Abstract

Diabetes is a global health problem that was estimated to be the 7^th^ leading cause of death worldwide in 2016. Type 2 diabetes mellitus (T2DM) is classically associated with genetic and environmental factors, however recent studies have demonstrated that the gut microbiome, which is altered in T2DM patients, is also likely to play a significant role in disease development. Despite this, the identity of microbiome-derived metabolites that influence T2DM onset and/or progression remain elusive. Here we demonstrate that a serum biomarker for T2DM, previously of unknown structure and origin, is actually two microbiome-derived metabolites, 3-methyl-4-(trimethylammonio)butanoate (3M-4-TMAB) and 4-(trimethylammonio)pentanoate (4-TMAP). These metabolites are produced by the *Lachnospiraceae* family of bacteria, which are highly prevalent in the gut microbiome of T2DM patients and are associated with high dietary fat intake. Treatment of human liver cells with 3M-4-TMAB and 4-TMAP results in a distinct change in the acylcarnitine profile in these cells and significantly reduced their insulin sensitivity; both indicators of T2DM. These results provide evidence of a mechanistic link between gut microbiome derived metabolites and T2DM.

## Main

Approximately 463 million people are living with diabetes worldwide and it is estimated that type 2 diabetes mellitus (T2DM) accounts for 90% of all cases^1^. Disruption of the gut microbiome is linked to various metabolic diseases and in T2DM it has been directly implicated in disease progression through alteration of insulin sensitivity^2^. *Lachnospiraceae* are anaerobic bacteria that are intrinsically linked to Western disease, with their presence in the human gut microbiome correlating with both a high fat diet and antibiotic use^3–5^. We recently discovered a mechanism by which two *Lachnospiraceae*-derived metabolites, both capable of crossing the blood brain barrier, directly inhibit energy production in human cells isolated from the white matter^6^. These metabolites, 3-methyl-4-(trimethylammonio)butanoate (3M-4-TMAB) and 4-(trimethylammonio)pentanoate (4-TMAP), are produced by the commensal strains *Clostridium clostridioforme* and *Clostridium symbiosum*. Increased numbers of these strains alongside other members of the *Lachnospiraceae* family are present in the gut microbiomes of obese individuals and in patients with T2DM^7–10^. The fact that 3M-4-TMAB and 4-TMAP have the capability to induce mitochondrial dysfunction and reduce functional capacity as observed in T2DM^11^, led us to hypothesize that they may have an important role in the development of T2DM. A role for the microbiome, and thus microbiome derived metabolites, in obesity and T2DM is further strengthened by evidence that germ free mice are protected from diet-induced obesity due to elevated rates of fatty acid oxidation (FAO) compared to colonised animals^12^ and that an obese phenotype can be induced in mouse models via microbiota transplantation alone^13,14^. Additionally, a metabolite(s) of the same mass as 3M-4-TMAB and 4-TMAP has been detected in patients undergoing treatment for T2DM and has also been described as a disease biomarker in a mouse model of T2DM^15,16^.

To determine any role for 3M-4-TMAB and 4-TMAP in T2DM, we first confirmed their presence in serum obtained from patients with T2DM and control patients. 3M-4-TMAB and 4-TMAP, which have the same mass, were detected in serum through mass spectrometry (Fig.1). The identification was validated by comparing the metabolites detected in the serum to metabolites detected in *C. symbiosum* lysates (previously shown to produce 3M-4TMAB and 4-TMAP) by tandem mass spectrometry (MS/MS)^6^. Fragmentation of the precursor ion (*m/z* 160.1323, [M+H]^+^ of 3M-4TMAB and 4-TMAP Δ −5.6 ppm) resulted in the same four fragment ions from both the metabolites in the serum and the *C. symbiosum* lysates. Furthermore, these four fragment ions matched those shown previously in the identification of 3M-4TMAB and 4-TMAP^6^. Detection of other peaks in the spectra were generated adjacent to the spotted samples and hence indicated they were not produced from the serum or bacteria samples and are likely background noise (Fig. 1c). Figure 1d shows the structures of 3M-4-TMAB and 4-TMAP and the chemical formulae of the assigned precursor and fragment ions. These results confirm that the metabolites from the patient serum and bacteria are the same, i.e. they both arise from 3M-4-TMAB and 4-TMAP.

**Figure 1.**
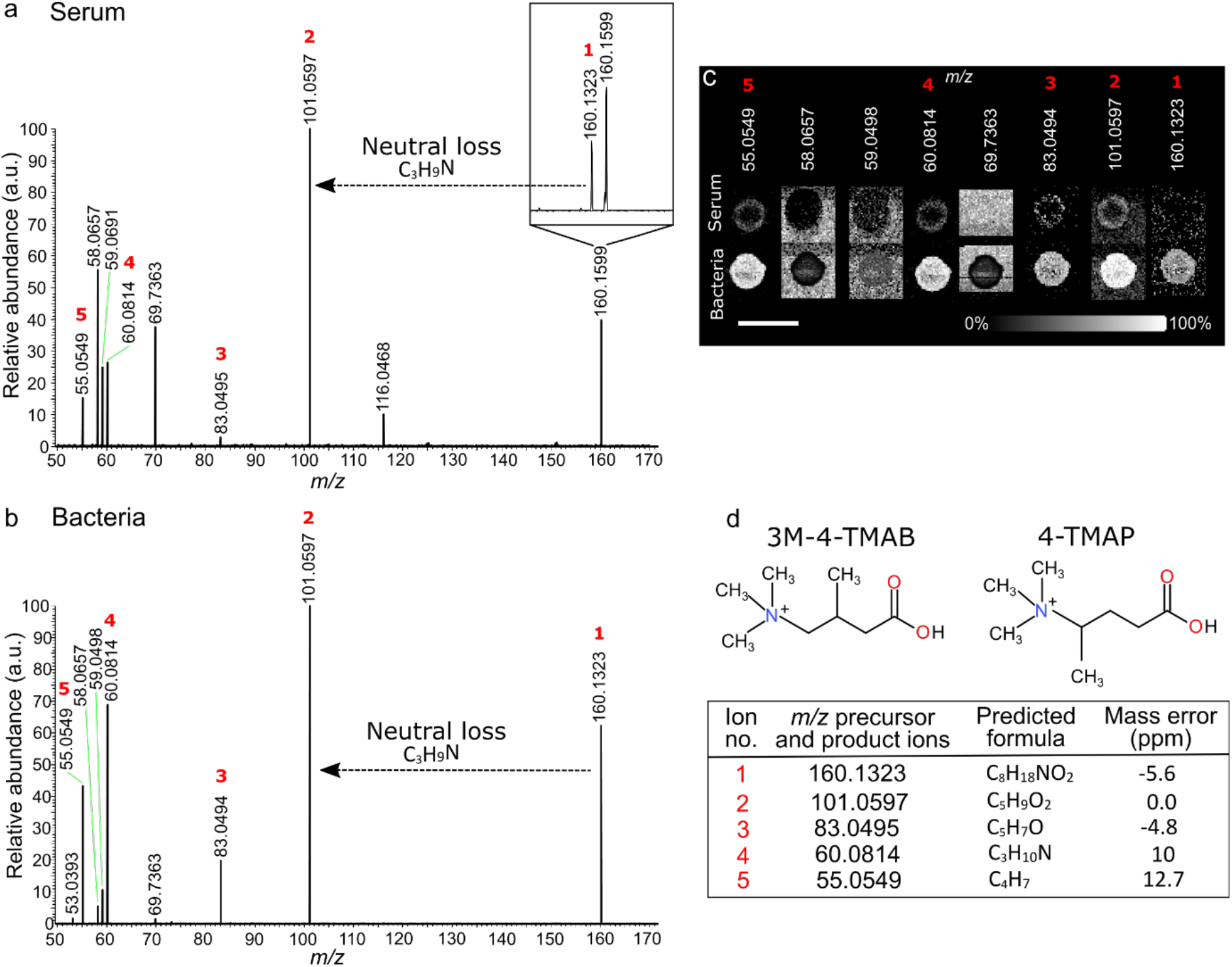
MS/MS analysis confirms serum metabolites are 3M-4-TMAB and 4-TMAP. MS/MS spectra from the fragmentation of the positive ion at *m/z* 160.1323 ([M+H]^+^ ion of 3M-4-TMAB and 4-TMAP) from diabetic patient serum (a) and bacteria (b). The fragment ions generated from dissociation of precursor ion at *m/z* 160.1323 are labelled with red numbers. MS/MS images of the ions in the spectra showing if the ion was generated from the bacteria and serum spot or from background noise (c). Scale bar = 5 mm. The structures of 3M-4-TMAB and 4-TMAP are shown with a table identifying the formula for each fragment ion from the MS/MS of *m/z* 160.1323 from the bacteria and serum (d). The neutral loss of C3H9N is also annotated in the spectra. Note the MS studies do not define the stereochemistry of 3M-4-TMAB and 4-TMAP.

Dysregulated FAO resulting in tissue lipid accumulation is associated with the development of insulin resistance^17^; these defects manifest as alterations in acylcarnitine abundance due to defective trafficking and breakdown. Therefore, we quantified acylcarnitine levels in human HepG2 liver cells after 72 hours of treatment with 3M-4-TMAB and 4-TMAP, relative to untreated cells. A significant decrease in total acylcarnitine levels was detected (Fig. 2). Additionally, decreased levels of specific medium chain acylcarnitines (e.g. 12:0 - dodecanoylcarnitine) and long chain acylcarnitines (e.g. 14:0 - tetradecanoylcarnitine, 14:1 - tetradecenoylcarnitine, 16:1 – hexadecenoylcarnitine, and 18:1 - oleylcarnitine) were detected in HepG2 cells in the presence of the bacterial metabolites (Fig. 2), a phenomenon which has previously been described as an indicator of T2DM in patients ^17^.

**Figure 2:**
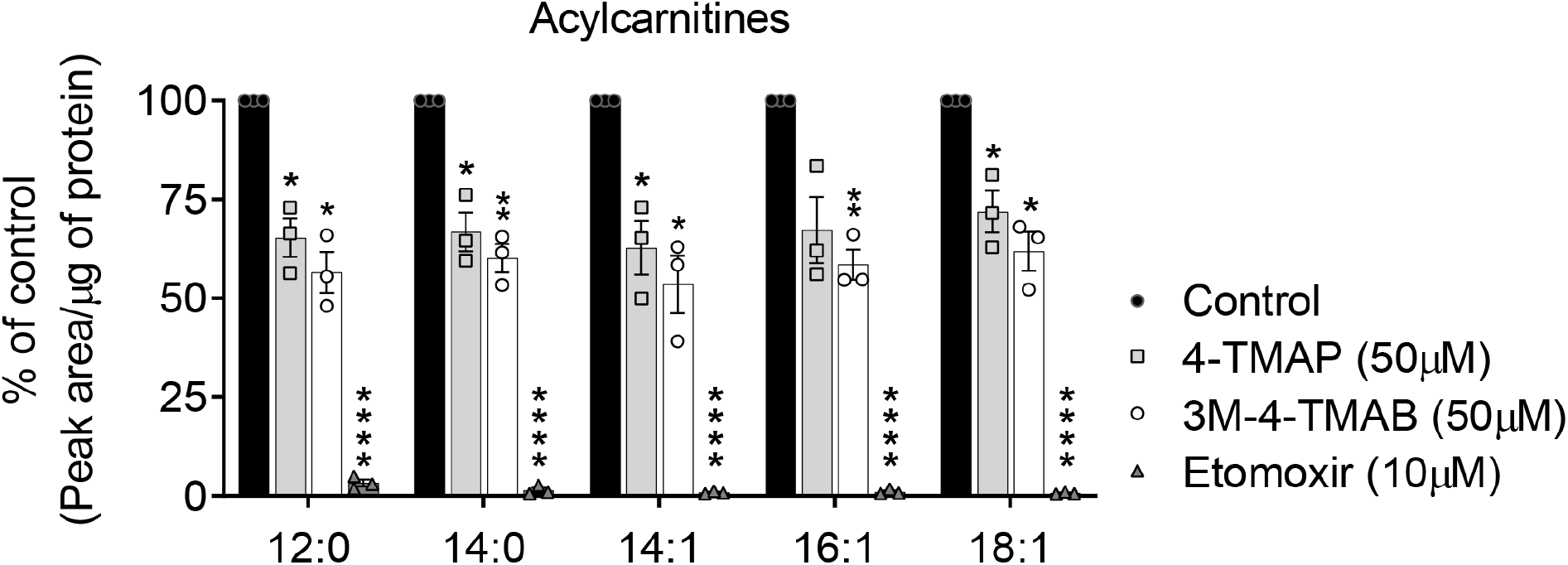
Acylcarnitine profiling of HepG2 cells after administration of the bacterial metabolites 4-TMAP and 3M-4-TMAB. HepG2 cells were treated with either 50 μM 4-TMAP or 3M-4-TMAB for 72h before acylcarnitine extraction and analysis. The effects of etomoxir (10 μM), an irreversible inhibitor of fatty acid oxidation, are shown for comparison. Results are presented as % of the values obtained for vehicle treated cells (Control). Bars represent mean ± SEM, symbols represent mean values from each independent experiment. Statistical significance was determined via a one sample t test (* = *P*< 0.05, ** = *P*< 0.01, ****= *P*<0.0001).

Having established that FAO inhibition occurs in HepG2 cells we then investigated the impact that 3M-4-TMAB, the more effective of the two metabolites in reducing acylcarnitine levels, had on insulin resistance in these cells. Insulin resistance leading to an accumulation of glucose in the bloodstream is a major contributory factor in T2DM and can be induced by increased hepatic fatty acid/triglyceride synthesis and reduced FAO. The metabolic actions of insulin are mediated via signalling pathways that lead to increased activation of Akt via phosphorylation at Ser473 and Thr308^18^. We therefore assessed phosphorylation of Akt at Ser473 phosphorylation following insulin-stimulation in HepG2 cells preincubated with 3M-4-TMAB. Preincubation of HepG2 cells with 3M-4-TMAB led to a significant reduction in phosphorylation of Akt at Ser473 upon exposure to increasing concentrations of insulin, indicating a reduced sensitivity to insulin (Fig. 3). This demonstrates that 3M-4-TMAB is capable of inducing mitochondrial dysfunction and insulin resistance in human liver cells; both characteristic phenotypes of T2DM.

**Figure 3:**
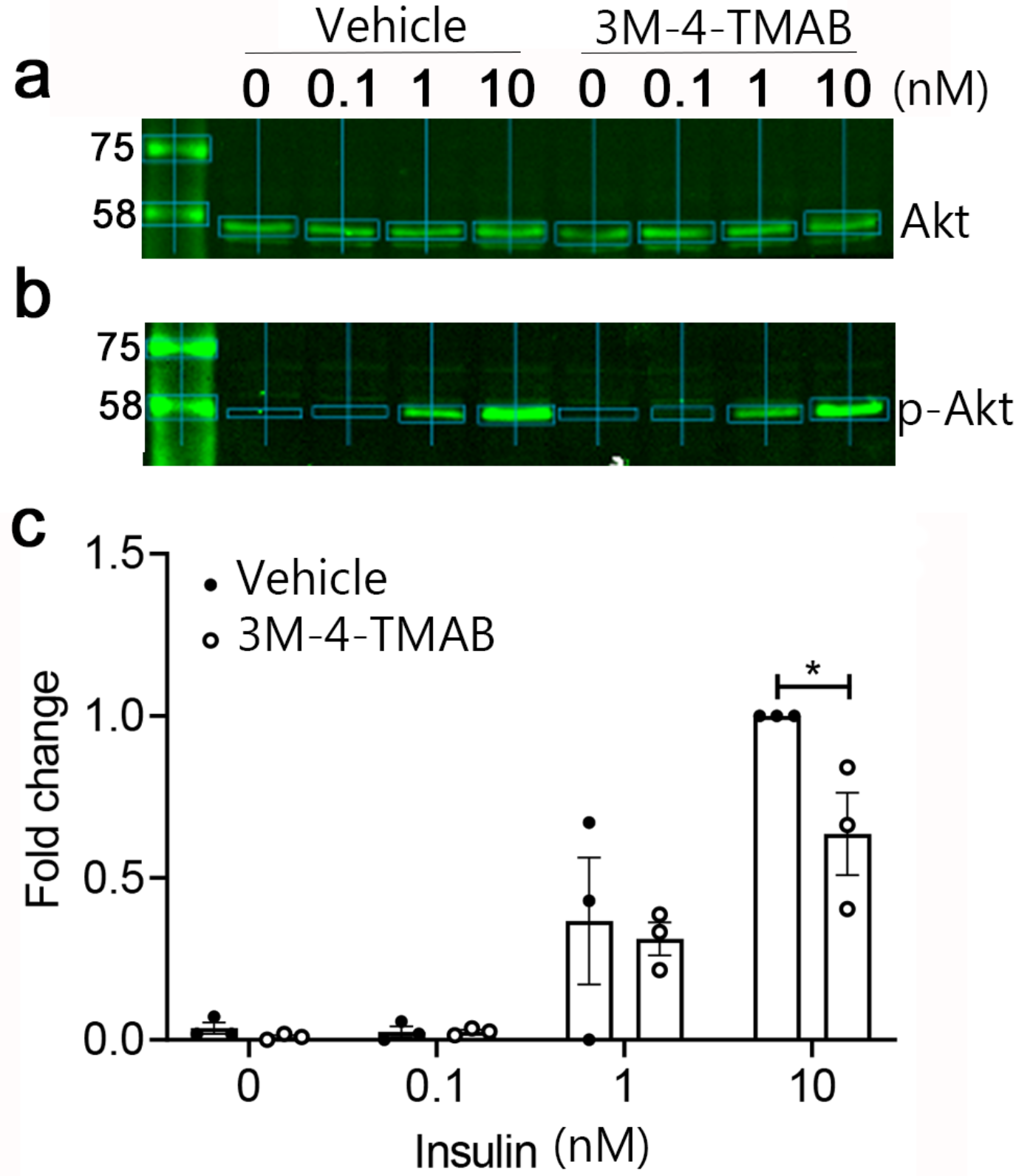
Pre-incubation of HepG2 cells with 3M-4-TMAB significantly reduces insulin-stimulated Akt Ser473 phosphorylation. Insulin-stimulated Akt Ser473 phosphorylation was assessed in HepG2 cells preincubated with 3M-4-TMAB (50 μM) for 72 hours. Insulin stimulation was carried out using the concentrations indicated (nM) for 1 hour prior to cell lysis. Total Akt (**a**) and Akt Ser473 phosphorylation (**b**) were detected by Western blotting of cell lysates. (**c**) Quantification of fold change in Akt Ser473 phosphorylation relative to total Akt levels (mean ± SD). Statistical significance was determined via a two-way ANOVA (* = *P*< 0.05).

Our results provide mechanistic evidence that the bacterially derived molecules, 3M-4-TMAB and 4-TMAP, which are detected systemically in animal models and in human serum, contribute to disease pathogenesis in T2DM. To our knowledge this is the first molecular evidence directly linking the gut microbiome to T2DM. The systemic presence of the metabolites and their ability to significantly impair mitochondrial function in human liver cells, and in cells isolated from the murine white matter^6^, leads us to hypothesize that they have wide-ranging implications for mammalian physiology. We propose that an increase in systemic levels of 3M-4-TMAB and 4-TMAP may be a contributing factor in a wide range of diseases where disruption of the gut microbiome disruption and mitochondrial dysfunction of an unknown cause are key clinical features.

## Methods

### Bacteria and growth conditions

Bacterial strains *Clostridium symbiosum* LM19B, isolated as previously described^6^, were routinely cultured on fastidious anaerobe broth (FAB) agar plates at 37°C under anaerobic growth conditions. For mass spectrometry imaging (MSI) analysis, single colonies from freshly cultured FAB agar plates were inoculated into 100 μl of phosphate buffered saline (PBS) to give an optical density at 600 nm of 0.3. Each bacterial culture (2 μl) was spotted onto a poly-L-lysine (in boric acid buffer [0.1 mg/ml; pH 8.4]) coated glass slide and allowed to dry at room temperature. Slides were stored in a desiccator until MS/MS was conducted as described below.

### Patient samples

Stored (−80°C) serum samples from participants of previous studies, VAPOUR (ClinicalTrials.gov identifier NCT03358953) and PRIORITY (ClinicalTrials.gov identifier NCT02040441), were retrieved for MS/MS analysis. They included 10 randomly selected samples from patients with T2DM (50% female; mean age 56.9 years) and 10 randomly selected samples from study participants with no history of overt cardiovascular disease (30% female; mean age 43.8 years). All participants provided written informed consent for future use of stored samples. These studies adhere to the principles of the Declaration of Helsinki and were approved by the West of Scotland Research Ethics Committee (reference 13/WS/0284) and the East of Scotland Research Ethics Committee (reference 16/ES/0103).

### Tandem Mass Spectrometry

MS/MS analyses were carried out using an Orbitrap mass spectrometer (Q Exactive, Thermo Fisher Scientific) equipped with a Prosolia 2D DESI source (Prosolia OmniSpray 2D). The source was modified with the spray tip positioned 1.5 mm above the sample surface and an angle of 75°. There was a 7 mm distance between the sprayer and the mass spectrometer inlet and a collection angle of 10°. The spray solvent was methanol/water (95:5, v/v), delivered at 1.5 μl/min using a Dionex Ultimate 3000 pump (Thermo Fisher Scientific). The Q Exactive settings were: 50 V S-Lens, positive ion mode, 50-170 *m/z* range, 1000 ms injection time for the individual MS/MS spectra, 500 ms injection time for the MS/MS imaging experiment, and a mass resolution of 70,000. MS/MS was performed using high collision dissociation with a normalized collision energy of 50 % for the individual spectra and 65 % for the imaging experiment. The spatial resolution was set at 150 μm.

### Hep-G2 cell culture

Hep-G2 cells were obtained from the American type culture collection (ATCC; HB-8065) and were cultured in Eagle’s Minimum Essential Medium (EMEM; Sigma) supplemented with 10% Foetal bovine serum (FBS; Invitrogen), 2 mM L-glutamine (Sigma) and 100 U/ml Penicillin/Streptomycin (Sigma). Cells were seeded at a density of 1 × 10^5^ cells per well in a 12-well plate and allowed to reach confluency. One day prior to experiment, media was replaced with low serum media containing 3% FBS.

### Insulin sensitivity

Fifty micromolar 3M-4-TMAB was added to appropriate wells of Hep-G2 cells, cultured as described above, for 72 hours. Ten minutes prior to harvest, insulin was added at concentrations of 0 nm, 0.1 nm, 1 nm and 10 nm. Cells were then placed on ice and washed with phosphate buffered saline (PBS) prior to lysis in Triton X-100-based lysis buffer (50 mM Tris-HCl, pH 7.4 at 4°C, 50 mM NaF, 1 mM Na_4_P_2_O_7_, 1 mM ethylenediaminetetraacetic acid, 1 mM ethylene glycol tetraacetic acid, 1% (v/v), Triton-X-100, 250 mM mannitol, 1 mM DTT, 1 mM Na_3_VO_4_ and 1 complete mini protease tablet). Cell lysates were scraped into microcentrifuge tubes and incubated on ice for 20 minutes, centrifuged (5 minutes, 21910 g, 4°C) and the subsequent supernatants stored at −20°C. Lysate protein concentrations were determined using a BCA protein assay kit according to manufacturer’s instructions (ThermoFisher).

Cell lysate proteins were resolved by SDS-PAGE and immunoblotted with antibodies diluted in TBS containing 0.1% (v/v) Tween-20. Antibodies used were: Akt (pan) (40D4) Mouse mAb #2920; and Phospho-Akt (Ser473) Antibody #9271 (both Cell Signalling Technologies, UK). Proteins were visualised using infrared dye-labelled secondary antibodies on a LI-COR Odyssey infrared imaging system and analysed using Image Studio Lite and Empiria for densitometric quantification of band intensity. In all cases, immunoblots for phospho- and total protein levels of Akt were obtained on immunoblots conducted concurrently in parallel. Three independent biological replicates were conducted.

### Acylcarnitine profiling

For acylcarnitine profiling HepG2 cells were cultured Minimum Essential Medium (MEM, Gibco cat. N. 11090081) supplemented with 10% FBS, 0.65 mM *L*-glutamine, and 1% Gibco MEM non-essential amino acids. Cells were seeded at 3 × 10^5^ cells/well in a six well plate. The day after seeding the medium was replaced with Plasmax™,^19^ supplemented with 2.5% FBS, and 200 mg/L of Albumax− II lipid rich bovine serum albumin. The following day Plasmax™ was refreshed and cells were incubated with 0.5% water (vehicle control) or 4-TMAP (50 μM), 3M-4-TMAB (50 μM), etomoxir (10 μM) for 72 h. The cells were then washed two times with ice-cold PBS and extracted with 0.4 mL of a butanol:methanol (1:1) solution, supplemented with Splash Lipidomix (Avanti Polar Lipids, Alabaster, AL, USA) as an internal standard. Lipids were extracted at −20°C for 15 min, the extracts were collected in tubes and centrifuged (16000 g × 10 min at 4°C,); the supernatant was transferred to high performance liquid chromatography (HPLC) vials for MS analysis. The total amount of cellular proteins was quantified in each of the extracted wells with a modified Lowry assay. The extracted lipids were separated and analysed with a Ultimate 3000 HPLC system (Thermo Fisher Scientific, Waltham, MA, USA) coupled to a Q-Exactive Orbitrap mass spectrometer (Thermo Fisher Scientific, Waltham, MA, USA) as described^20^. Peak detection and integration from Raw data were processed using Compound Discoverer 3.0 (Thermo Fisher Scientific, Waltham, MA, USA) and converted to mgf format using MSConvert software. Files were searched against LipidBlast database using LipiDex software^21–23^. Peak areas were normalised on the total amount of cellular proteins determined from extracted cells.

### Statistical analysis

For analysis of acylcarnitines three independent experiments were undertaken with each biological replicate shown. For each of the conditions tested, the value of acylcarnitine from each treated condition was expressed as relative to the vehicle treated control, and the significance of the differences between conditions was assessed with a one sample t test (relative value = 1, Graph Pad Prism 8.1.2). Insulin sensitivity was determined in three independent replicates for each concentration tested through measurement of fold change in Akt Ser473 phosphorylation between vehicle and 3M-4-TMAB treated HepG2 cells. Significance was determined through a two-way ANOVA. *P* values ≤ 0.05 were considered significant for all statistical tests.

## Acknowledgements

This work was supported by a Biotechnology and Biological Sciences Research Council (BBSRC)–CASE studentship in part funded by AstraZeneca (to R.B., D.M.W., and R.J.A.G.), BBSRC grants BB/K008005/1 and BB/P003281/1 (to D.M.W.) and a Diabetes UK equipment grant (BDA11/0004309 to I.P.S.). S.T. was supported by Cancer Research UK Beatson Institute core funding (C596/A17196) and CRUK core group award (A23982). CD was supported by the British Heart Foundation (RE/13/5/30177 and RE/18/6/34217) and European Commission (“PRIORITY”; grant agreement 279277). CJS was supported by Cancer Research UK and the Wellcome Trust. We would like to thank the Core Services and Advanced Technologies at the Cancer Research UK Beatson Institute (C596/A17196), with particular thanks to the Metabolomics facility. We also thank Dr. Lynsey Meikle (Kendall Medical Writing) for editorial assistance in the preparation of this manuscript.

